# Genome dashboards: Framework and Examples

**DOI:** 10.1101/803403

**Authors:** Zilong Li, Ran Sun, Thomas C. Bishop

**Affiliations:** College of Engineering and Science, Louisiana Tech University, P.O. Box 10157 Ruston, LA 71272, USA

## Abstract

Genomics is a sequence-based informatics science and a 3D structure-based material science. Here we describe a framework for developing genome dashboards specifically designed to unify informatics with studies of chromatin structure and dynamics. The framework is based on the mathematical representation of geometrically exact rod models and the generalization of DNA base pair step parameters. A Model-View-Controller software design approach is proposed to implement genome dashboards as finite state machines either as desktop or web based applications. Two examples are demonstrated using our minimal genome dashboard called G-Dash-min. The data unification achieved with a genome dashboard supports the bi-directional exchange of data between informatics and structure. Thus any experimentally or theoretically determined sequence based informatics track can inform DNA, nucleosome or chromatin modeling (e.g. nucleosome positions) and structure features can be analyzed as informatics tracks in a genome browser (e.g. DNA base pair step parameters: Roll, Tilt, Twist). Here the framework is applied to chromatin, but genome dashboards are more broadly applicable. Genome dashboards are a novel means of investigating structure-function relationships for regions of the genome ranging from base pairs to entire chromosomes and for generating, validating, and testing mechanistic hypotheses.

## INTRODUCTION

Chromatin is the biomaterial that contains the genome in all higher organisms. There is no consensus on the structure of chromatin (1), but there is a wealth of informatics and structure data available. From an informatics point of view, efforts such as the 1000 Genomes (2), ENCODE (3) and the 4D Nucleome (4) projects provide sequence based reference data. Coupling the reference data with Next Generation Sequencing (NGS) and informatics analysis pipelines enables individual labs to conduct Genome Wide Association (GWA) studies that link chromatin reprogramming with disease and altered gene expression as described, for example, in (6, 7).

Hi-C (8, 9), Micro-C (10) and other chromosome conformation capture methods (11, 12) provide distance constraints as a measure of the large scale organization of chromatin structure. Super resolution microscopy (13, 14) provides optical visualization of 3D structure at nanometer scale resolution, while electron microscopy (15), NMR (16), and X-ray crystallography (17) provide ångström scale resolution of individual nucleosomes and nucleosome arrays. Strategies for modeling chromatin structures are rapidly maturing (18, 19, 20). The desire to merge computational and experimental approaches is recognized (21, 22), but a significant challenge in chromatin structural biology is unifying these diverse data sets to advance our understanding of structure-function relationships and to validate genomic mechanisms of action.

Here we label sequence based informatics as a 1D representation, contact and distant constraints as 2D representations, X-ray, NMR and super-resolution microscopy as 3D representations and dynamics as a 4D representation of chromatin. There exists a growing collection of computational tools that convert 2D data to 3D structures of chromatin (https://www.4dnucleome.org/software.html), and computational models can promote 3D structures to dynamics or sampling data (4D). But, there remain few tools, other than ICM-Web (23), that directly link sequence (1D) with chromatin structure (3D) or dynamics (4D). Thus researchers utilizing 1D sequence based methods are missing 3D structural data including steric and geometric constraints in their analyses, and researchers utilizing 3D and 4D computational and experimental methods are missing the wealth of informatics data available in sequence based data sets.

A genome dashboard, like an automobile dashboard or airplane cockpit, integrates a console for managing information with controllers for navigating a physical world that appears in a window. A “genome dashboard” unifies informatics (1D), contact and two-angle representations (2D) (24), structure (3D), and dynamics (4D) data describing DNA, nucleosomes and chromatin. For the purpose of developing such genome dashboards, we have identified a framework that unifies 1D and 3D representations and a general method for implementing it that supports data visualization and manipulation. The framework is bi-directional, i.e. can map 3D representations to 1D and 1D representations to 3D.

The framework is based on mathematical representations of geometrically exact rods (25), presented below as “The Model”, followed by “Design Considerations” for implementing it using Model-View-Controller software development principles. This approach enables a commodity-off-the-shelf (COTS) approach for assembling genome dashboards that is both extensible and portable. Two examples are then presented to demonstrate that G-Dash-min, a minimal web based implementation of a genome dashboard, can function in real time in both directions to convert informatics data into a physical model and a physical model into informatics data.

## FRAMEWORK

### The Model

From the genome dashboard perspective, informatics (1D) is any data that maps to DNA sequence. Generally speaking, NGS relies on aligning experimental data with chromosome coordinates and information theory for analysis. Physical models are 3D/4D descriptions that employ energy functions and physical laws for analysis. The energy functions are typically grouped into external and internal energies. *U* = *U*_*ext*_ +*U*_*int*_, where *U*_*ext*_ captures through space interactions and *U*_*int*_ captures local conformation and dynamical properties. Both require knowledge of material properties and geometry. Material properties such as van der Waals radii, dielectric properties, partial charge distributions, moments of inertia, mass, stiffness parameters, bond angles, etc.. are all parameters associated with a specific physical model or force field. The model may employ atomic, coarse-grained or continuous medium approximations. Geometry is the structure of the physical model, it may be expressed in a laboratory (external) or material (internal) reference frame and may be obtained from theoretical or experimental techniques.

Our strategy for unification (merging data from different sources) is based on the idea that DNA is the common thread in chromatin structural biology. Unification is achieved through laboratory (Cartesian coordinate) and material (internal coordinate) representations of DNA as an oriented space curve or ribbon, Figure 1. Since unification is based on geometric considerations, our strategy is independent of the parameters associated with a specific physical model. Of course, associating an energy landscape with a physical model requires one to choose a physical model. Our framework does not provide this model but it does not restrict the user’s choice of model. Implementations of the genome dashboard concept may support only one or many models.

**Figure 1.**
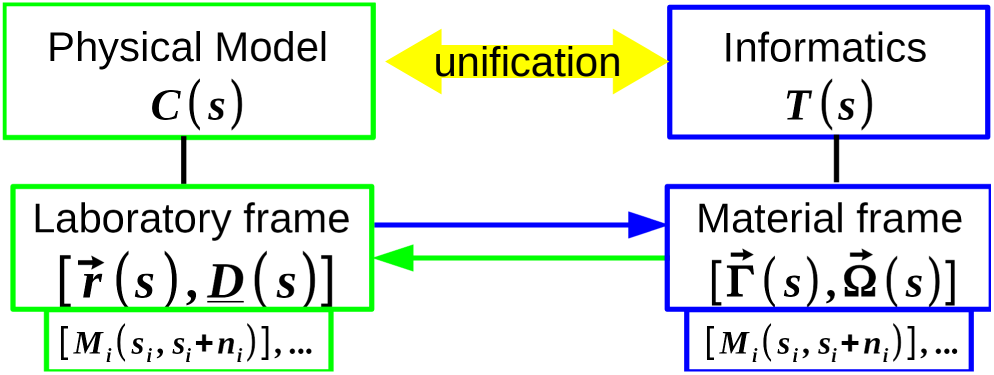
Unification is the process of merging data from different sources. Physical models and informatics data are unified by mathematical representations of an oriented space curve in laboratory 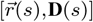 and material 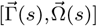 reference frames. The conformation of a physical model *C*(*s*) is associated with the laboratory frame, and informatics data *T*(*s*) is associated with the material frame. Masks *M*(*s*) alter the material properties of DNA. Exchanging data between laboratory and material frames unifies the physical model and informatics.

In a laboratory reference frame, a continuous ribbon has centerline 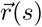 and unit length directors 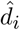 embedded in the ribbon that capture the local orientation of the ribbon. The directors can be represented by a director frame matrix 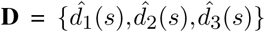 (26). This matrix also serves to transform a representation in the material(internal) frame to a representation in the laboratory (Cartesian) frame.

An equivalent description of the ribbon is based on the director frames themselves. This description is a material reference frame description that captures the translations and rotations connecting one director frame to the next, represented here by 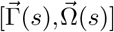. The two representations 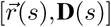 and 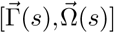 are equivalent descriptions of the conformation of an oriented space curve, denoted simply as *C*(*s*) for the Cartesian Coordinate representation and *T*(*s*) when expressed in the material frame and interpreted as informatics tracks.

Converting between the 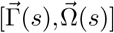 and 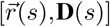 representations requires either a differentiation or an integration as expressed by the following equations.

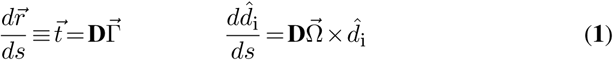

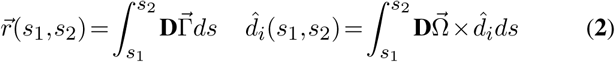

Here 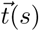 is the unnormalized tangent to the ribbon expressed in the laboratory frame. 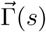 is the same vector expressed in the internal frame. 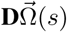 is recognized as the vector corresponding to the instantaneous axis of rotation of the director frames located along the ribbon at position *s*, as represented in the laboratory coordinate frame. Discrete approximations to Equations **1** and piecewise integration as expressed in Equations **2** can be employed to obtain a collection of discrete director frames. Different models may require different numerical algorithms to achieve the required discretization. Numerical methods suitable for DNA are discussed below. They should be reading strand invariant. Together, the two representations provide a basis for unifying informatics 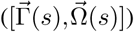 and structure 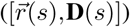 data.

DNA conformation *C*(*s*) is at best a base pair discrete approximation to a continuous oriented space curve (27, 28). Base pair step parameters (29, 30) and associated algorithms provide established methods for describing double and single stranded DNA as a discrete oriented space curve at atomic resolution. A sequence specific di-nucleotide accurate model of dsDNA in the 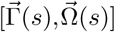 (base pair step parameter) representation can be obtained from x-ray (31) or molecular dynamics (32, 33) studies. A 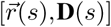 description is obtained by integrating 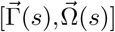. There are two widely used tools for base pair step parameter analysis. 3DNA (34) uses a Euler Angle (E-A) based method and employs a “RollTilt” approximation (35). Curves+ (36) uses a Euler-Rodrigues (E-R) based method (37). Both methods utilize a mid-step plane construction to ensure that the computed parameter values are not affected by the choice of reading strand. Mathematically, one must invert the sign of Tilt and Shift upon strand reversal to preserve the alignment of director frames with the DNA major and minor groove (38).

Helical parameter values obtained from the 3DNA and Curves+ methods are known to differ (39). Differences may arise from two sources. The assignment of director frames to base pairs may differ. However, the methods for assigning director frames are well-defined for ideal pairing (30), so these differences typically occur only for significant deviations from ideal geometries. The other source of differences is method dependent. These differences have not been well studied. Helical parameter values obtained from 3DNA and from Curves+ differ even when the director frames used for the calculations are identical, i.e. even when the first problem is eliminated. Thus, values obtained from one method should not in general be interchanged with the other. Nonetheless these two methods work well for all-atom representations of the pairing and stacking of base pairs in double stranded DNA with the caveat that neither provides information about the DNA backbone. Recent efforts now support proper reconstruction of the DNA backbone (40).

Chromatin, for our purposes, is a biomolecular structure composed of DNA and external agents, such as histones, that alter the material properties (conformation, dynamics, flexibility, energetics, chemical properties) of a contiguous length of DNA from *s* to *s*+*n*. We label any such external agent, including histones, as a Mask, *M* (*s,s*+ *n*). Any number of identical or unique Masks (*M*_*i*_(*s*_*i*_,*s*_*i*_ + *n*_*i*_)) may be associated with a sequence of DNA (*s*_*i*_ to *s*_*i*_ +*n*_*i*_). With this approach, chromatin folding is the process of activating Masks. There are two strategies for activating Masks. The first is achieved in the material reference frame with the conformation of the masked DNA denoted by a list of internal coordinates 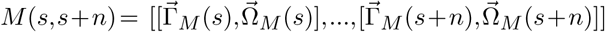 that spans *n* base pairs. As the name suggests, the Mask replaces the 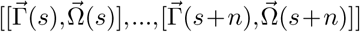 values associated with DNA. We can calculate *C*(*s*) from Equations **2**, as discussed above. The second approach is achieved in the laboratory reference frame with the conformation of DNA described as a rigid entity with 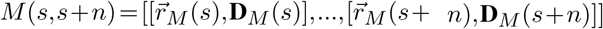. The Mask consists of Cartesian coordinates and director frames which can be converted to *T* (*s*) using Equations **1** as discussed above. The Cartesian coordinate representation of *M* (*s,s*+*n*) requires only a single translation and rotation to position each rigidly masked element in the laboratory reference frame.

In terms of Masks, a nucleosome is a DNA superhelix *and* histones. The histones can be represented independently of DNA as a single entity (sphere, cylinder, ellipsoid), a collection of beads, or an all-atom model. Alternatively the DNA *and* histones can be included in the nucleosome Mask as a single entity. Docking individual histones or the complete histone octamer to a superhelix or placing entire nucleosomes between linkers can be achieved with the same methods and tools used for describing the relative rotations and translations of base pairs. However the “RollTilt” small angle approximation is no longer valid. Linker DNA connecting Masks is often assumed to be free DNA, but in general, even linker DNA may be described by Masks, e.g. bent or less flexible linkers. Likewise chemical modification of the DNA, e.g. methylation, which does not change the sequence but the physical-chemical properties of DNA, is a Mask.

In the context of a genome dashboard, chromatin folding is an informatics problem of describing all the unique Masks and tracking their locations along a sequence of DNA. These Masks can be manipulated based on informatics or physical analyses. In this manner, genome dashboards are designed to enable users to efficiently define and navigate chromatin folding landscapes.

### Design Considerations

A genome dashboard is a finite state machine that can be efficiently developed using Model-View-Controller (MVC) design principles (41), Figure 2. This approach ensures that a dashboard’s components are independent, replaceable, and extensible.

**Figure 2.**
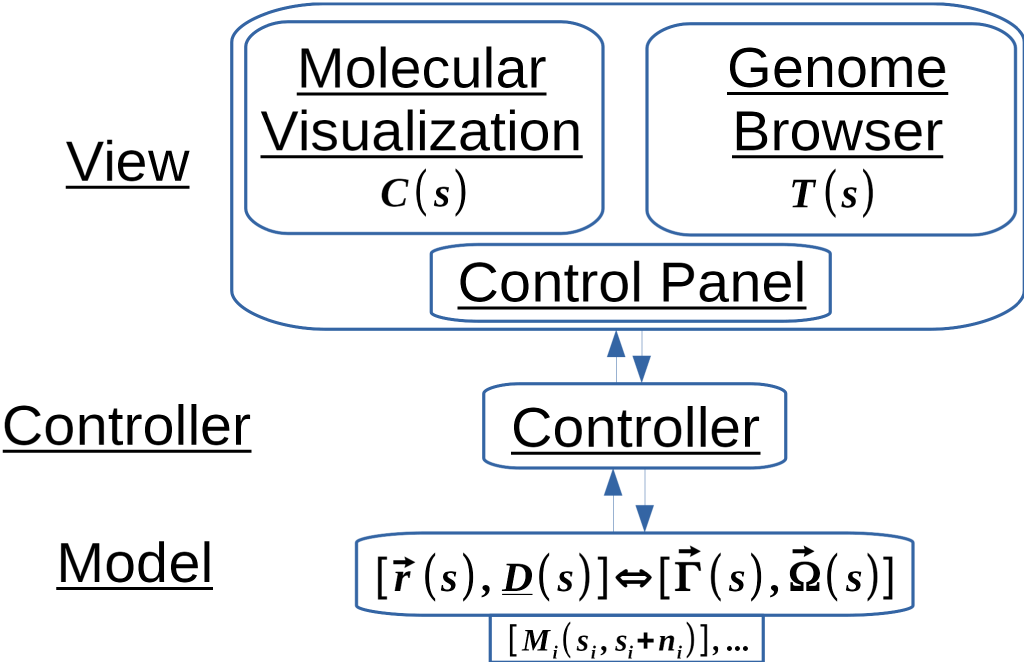
Model-View-Controller (MVC) Design. Model: Laboratory frame 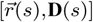 and material frame 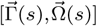 descriptions of DNA as the common thread, an inventory of Masks *M* (*s*), and procedures for converting between representations. View: a Molecular Visualization (MV) displays *C*(*s*), a Genome Browser (GB) displays *T*(*s*) and a Control Panel (CP) provides a graphical interface to the controller. G-Dash-min uses JSmol and Biodalliance for the MV and GB components, respectively. A commodity-off-the-shelf (COTS) approach enables a genome dashboard to use any desired MVs and GBs. Controller: manages the exchange of data between Model and Views.

The “Model” in the MVC schema is the data and related logic. For a genome dashboard, the Model includes the 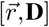 and 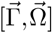 representations of DNA as a discrete (or even continuous (42, 43)) oriented space curve, the inventory of Masks, *M*_*i*_ (*s*_*i*_,*s*_*i*_ +*n*_*i*_), any associated track data, *T*(*s*), and procedures for converting between representations. In general, the rotations and translations associated with a Mask may be large. If a Mask is a rigid entity, this information can be leveraged to improve performance. For example, representing all 147 base pairs of DNA and eight histones in a nucleosome as a single director frame along with a large deformation of the path of DNA reduces computational and data costs by approximately *n* * 146, where *n* is the number of nucleosomes containing 147 base pairs.

The “View” in the MVC schema provides the user interface and renders data. A genome dashboard includes a 3D Molecular Visualization (MV) for rendering 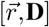, a Genome Browser (GB) for rendering 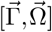, and a Control Panel (CP) as a graphical user interface to the Controller. A genome dashboard can be designed as a web application or a stand alone application. For web applications, javascript based MVs such as JSmol (44) and NGL Viewer (45) are optimal. For stand alone applications, MVs such as VMD (46) and PyMOL (The PyMOL Molecular Graphics System, Version 2.0 Schrödinger, LLC.) are optimal. Likewise, for web applications, the GB should be javascript based, like Biodalliance (47). For stand alone applications, JBrowse (48) or other modern genome browsers may provide advantages.

The “Controller” in the MVC schema manages the exchange of data between the View and the Model. For a given genome dashboard, the MV and GB can be COTS elements, but the Control Panel and Controller are application specific. We expect that different instances of the genome dashboard concept will target different users and utilize different physical models. Different strategies for managing the MV, GB, and CP may emerge, but the underlying Model remains as described above.

## EXAMPLES

Based on the framework proposed above, we have developed a minimal genome dashboard named “G-Dash-min”. Here we demonstrate two examples using G-Dash-min to show how genome dashboards contribute to our understanding of biological function by the unification of informatics and physical models.

### Informatics to physical model

A hormone response element (HRE) is a specific sequence of DNA representing 15 base pairs. Selective binding of an activated hormone receptor (HR) to the HRE is a critical component of the hormone response mechanism, see Fig 1-41 of (49). A variant of this gene regulatory mechanism is employed to control numerous physiologic functions in all higher organisms. To demonstrate the power of data unification achieved with our G-Dash-min application, we have identified estrogen response elements (ERE) using ERE-Finder (50). This informatics data is displayed as the ERE Track in Fig 3. All experimentally determined nucleosome positions for the human genome (51) are also displayed in Fig 3 as the Nuc-Pos Track. These Tracks provide locations for EREs and nucleosomes; however, the 1D representations are insufficient to determine whether or not the locations are physically realizable. These tracks are sufficient to identify several regions of interest. One of them is associated with chromosome 6 and coordinate location 168,131,722 to 168,132,130. Here we find three overlapping nucleosomes and an ERE that appears to function as a classic switching mechanism. We explore this hypothesis with G-Dash-min by generating physical models.

**Figure 3.**
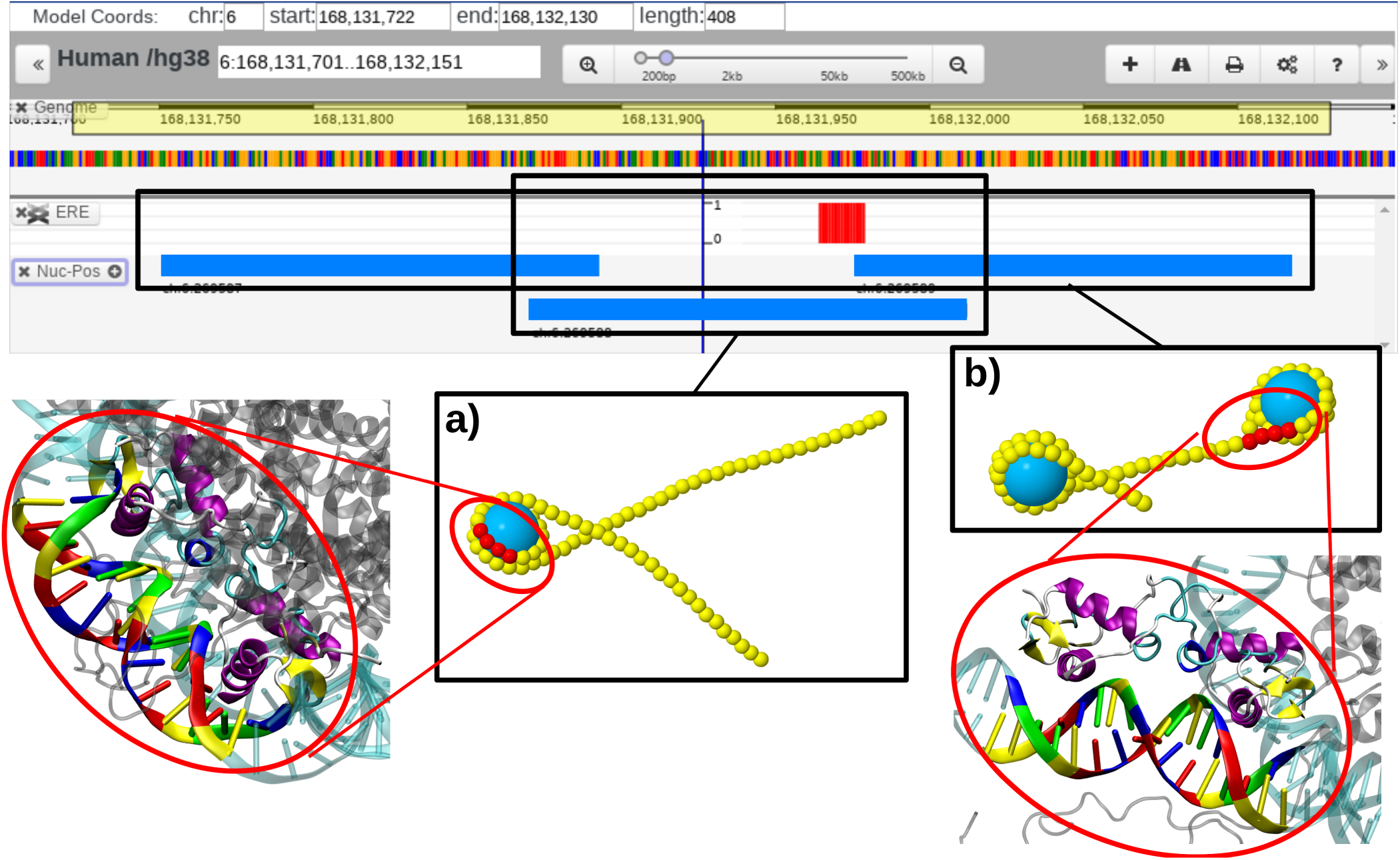
Black boxes: *C*(*s*) and *T*(*s*) representations of two allowed states. Upper boxes are *T*(*s*) representations of nucleosome positions (blue bars) and an ERE (red bar). Lower boxes are *C*(*s*) representations (small beads represent 5 base pairs large beads represent histone octamers). Red ellipses: The corresponding all-atom structures with the estrogen receptor DNA-binding domain docked to the DNA as in PDB entry 1HCQ. a) The ERE is located within a nucleosome with the major groove facing inward. The receptor is prohibited from binding. b). The ERE is located in a nucleosome free region. Docking 1HCQ indicates that the ERE is physically accessible.

We first selected the single nucleosome shown in the bottom of the Nuc-Pos Track in Fig 3 and generated a coarse-grained model. Mapping the ERE location onto the physical model provided its physical location, but without knowledge of major groove orientation one can still not determine the accessibility of the ERE site for ER binding. We then constructed an all-atom model. With the all-atom model, we see that the major groove is actually facing towards the histones. This prevents the estrogen receptor DNA-binding domain (1HCQ) from binding to this region of the DNA major groove. To bind 1HCQ to the all-atom model we downloaded the all-atom model from G-Dash-min then loaded it and 1HCQ into VMD. A simple VMD script fit the DNA in 1HCQ to the DNA in the G-Dash-min all-atom model. (This script and the models are provided as supplementary data.)

We then used the same approach to model the two nucleosomes shown in the top of the Nuc-Pos Track in Fig 3. Mapping the ERE location to the coarse-grained model suggests the ERE may be accessible to ER. Using the same procedure and script as before, we find that 1HCQ can physically access this ERE with the nucleosome present. We point out that 1HCQ is only the DNA binding domain of the estrogen receptor. There exist steric conflicts between the ER-DBD and the histones so this is not the complete story; but it strongly suggests this site as a candidate for a genetic switching mechanism.

With this example we demonstrate with G-Dash-min how informatics is used to construct a physical model that extends and validates the interpretation of the informatics data. We emphasize that any informatics track or combination of tracks can be used to inform the physical model.

### Physical model to informatics

Models of chromatin are rapidly maturing. As the models develop, there is increasing demand to capture biologic realism, including DNA sequence, experimentally determined nucleosome positions, states of chemical modification etc… Manually curating informatics data to build a “biologically inspired” model is a time-consuming and tedious task that informs the initial model but does not necessarily support the interpretation of modeling results. There exists a growing collection of published physical models to which such interpretations could be applied. Here we import a HOXC Mesoscale model generated by the Schlick lab (52) into G-Dash-min to demonstrate how informatics can be overlayed onto physical models to extend the interpretation.

In G-Dash-min we provide an upload function that is specific for DiscoTech based models (53). We upload and convert the HOXC Mesoscale model into a G-Dash-min model, Figure 4. The HOXC model utilizes a 9 base pair per bead model that includes both the location and orientation of each bead and each DiscoTech based nucleosome, i.e. 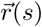 and **D**(*s*) are provided in the model. We treat the DiscoTech nucleosomes as Masks and calculate 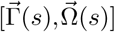 based on the 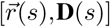 data provided for the beads and Masks. The 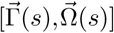 values computed are no longer DNA base pair step parameters; however, they still represent an oriented space curve or ribbon. We refer to them as generalized step parameters (GSP) and utilize the same naming conventions as for the DNA parameters: Tilt, Roll, Twist, Shift, Slide and Rise, and display them as informatics tracks in the genome browser. Our generalized parameter values differ significantly from those associated with DNA. Small angle approximations are no longer valid, see data in Figure 4-d.

**Figure 4.**
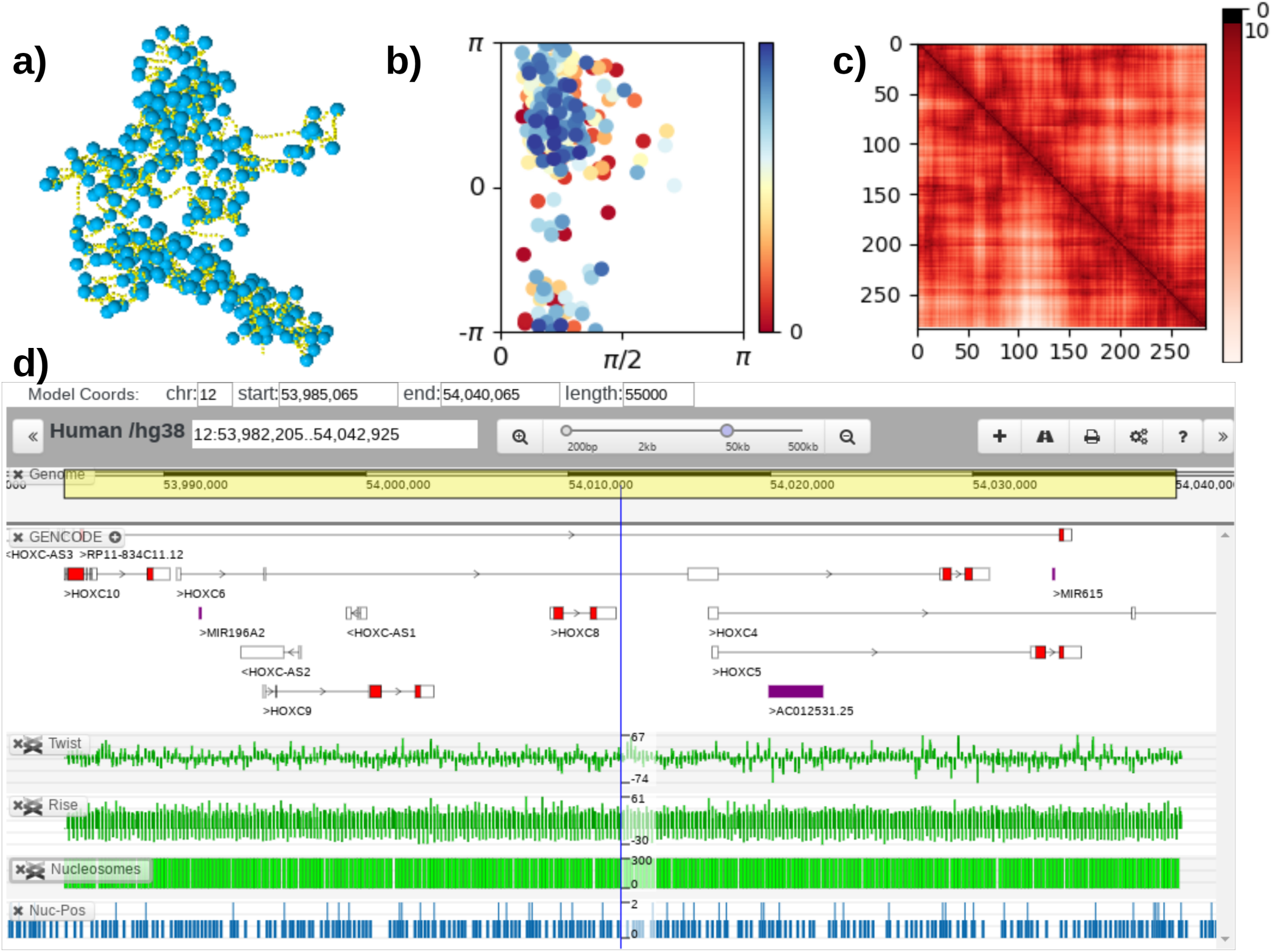
a) HOXC coarse-grained model of chromatin containing approximately 55k base pairs of DNA and 284 nucleosomes. Uploading the HOXC model to G-Dash-min generates: b) a two angle representation of the HOXC model, c) a distance-distance matrix based on nucleosome centers of mass, and d) structural informatics data. “Generalized Helical Parameter” (“Twist” and “Rise”) and nucleosome position (“Nucleosomes”) data are displayed along side experimentally determined nucleosome positions (“Nuc-Pos”) and other informatics data(“Gencode”).

To determine if either of the E-R or E-A approaches is suitable for our GSPs, we have converted all six HOXC models reported in (52) from the *C*(*s*) to the *T*(*s*) and back to the *C*(*s*) representations. Each G-Dash-min converted model contains approximately 1779 beads, represents 55,000 base pairs and has a unique conformation. The RMSD between the initial and final *C*(*s*) structures are computed using VMD’s RMSD functions. The E-R approach yields RMSD values ranging from 0.03*Å* to 0.15*Å* after alignment, and the E-A approach yields RMSD values ranging from 0.002*Å* to 0.003*Å* after alignment. Our experience is that both the E-A and E-R methods are acceptable algorithms for implementing “The Model” even when thermal fluctuations or deformations associated with nucleosomes and chromatin, or even generalized step parameters, are modeled. (All of these models are provided as supplementary data.)

The HOXC model was constructed for a specific sequence of DNA: HOXC10 of annotated human genome assembly 38 that begins at chr12:53,985,065. However, the data provided does not contain sequence information because the model itself is sequence independent. Thus, whenever a DiscoTech model is uploaded into G-Dash-min it must be aligned with sequence using the yellow bar sequence selection bar, Figure 4. If sequence information is included in the model, the model can be automatically aligned to data in the genome browser. Once aligned, the structural informatics tracks enable us to compare the nucleosome positions used in the model (Nucleosomes Track in Figure 4) to experimentally determined nucleosome positions (Nuc-Pos Track in Figure 4). It is clear that the two tracks differ. Resolving these differences promises to advance our understanding of both the experimental and modeling data.

To complete our multi-dimensional representation of chromatin folding, we have incorporated 2D representations into G-Dash-min. Figure 4-b is a two angle plot and Figure 4-c is a distance-distance matrix plot. As with the structure tracks, these are automatically generated for models containing sufficiently many nucleosomes. The two angle plot is a Woodcock Equivalent (WE) Plot (24, 54). On these plots, *α* is the angle between the centers of three adjacent nucleosomes, and *β* is the dihedral rotation angle obtained from the centers of four adjacent nucleosomes. For this analysis, the reported *α* and *β* values are computed as if the linkers were straight, even if the linkers are not. For this reason, we have adopted the label “Woodcock Equivalent” Plot. The distance-distance matrix reports the center-to-center distance between all nucleosomes. Unlike the exchange of data between the informatics (1D) and physical models (3D), the WE Plot data and distance-distance maps are one directional. The 2D representations are obtained from the 3D model but cannot be used to generate 3D models. Methods exist for this purpose but have not yet been implemented in G-Dash-min.

## DISCUSSION

As a working example of a genome dashboard, G-Dash-min demonstrates that informatics and physical models can be unified in a web based application in real time. Our tests demonstrate that interactive usage can be achieved for systems containing 10,000 to 50,000 base pairs or coarse-grained beads. The algorithms for converting informatics to physical models and physical models to informatics work in both directions for base pair resolution models using either the Euler-Rodrigues (Curves+) or Euler-Angle (3DNA) based method. We believe our application to the HOXC model is the first demonstration that both the Euler-Rodrigues and the Euler-Angle algorithms can be applied to coarse-grained models discretized well beyond the base pair level. We label the internal coordinates “generalized step parameters” in this case.

G-Dash-min also demonstrates that a commodity-off-the-shelf approach coupled with Model-View-Controller principles can be employed to efficiently develop genome dashboards that are customizable, extensible and portable. G-Dash-min generates atomic and coarse-grained models of DNA, nucleosomes, and chromatin using various experimental and theoretical data. Coupling G-Dash-min’s atomic modeling capabilities with high-performance, high-throughput workflows, and our TMB Library of nucleosome simulations (55) provides a software ecosystem for overnight comparative molecular dynamics simulations of nucleosomes (56). Such models are necessary for developing designer nucleosomes and assessing receptor-DNA interactions in their native context (57).

The genome dashboard framework achieves unification of informatics and physical models; the challenge is data representation. Genomics data has well-defined data formats, but there are numerous data formats for Cartesian coordinate data and no established conventions for representing director frame data or step parameters. We have demonstrated that the E-R and E-A algorithms can support multi-scale modeling of DNA as the common thread in chromatin using a geometrically exact rod model. However, we have deliberately avoided energetic considerations. The genome dashboard user or developer must decide which energy model is most appropriate for their particular application. We thus expect many genome dashboards to be developed, each tailored for a specific use.

As described here, genome dashboards are designed to work with chromatin folding, but the concept and framework are not limited to chromatin or even to eukaryotes. The informatics data can be any data associated with an indexing system, e.g. protein sequence. The structure data can be any slender polymer or potentially even an ordered collection of points in space that are linked to form an oriented space curve. The latter includes observations of nucleosomes with super-resolution microscopy. If the sequence ordering of nucleosomes in an image can be determined and director frames can be assigned to each nucleosome, it will be possible to unify 1D, 2D, 3D, and 4D representations of chromatin as in Figure 4.

## ACKNOWLEDGEMENTS

We thank Joohyun Kim and Jinghua Ge of the Center for Computation and Technology, Louisiana State University, and John Gentle of Science Gateways Community Institute (SGCI) for assistance and advice. We thank the Schlick Lab at NYU for sharing coordinate data for the HOXC models. Ran Sun’s contributions were related mostly to all-atom modeling and helix parameter analysis. Zilong Li developed G-Dash-min as a web-based tool and prepared the manuscript.

