## Supplemental Table 1 for "Genome dashboards: Framework and Examples"

### SUPPORTING MATERIAL

- G-Dash-min is available as a web application and as a Virtual Machine at the following url:
  http://dna.engr.latech.edu/~gdash/

- PDB files representing the binding of the ER-DBD to DNA as displayed in Figure 3 are included in ERE.tar.gz. The files included are: bottom.pdb and top.pdb corresponding to the two models shown the manuscript.

- The G-Dash-min HOXC models converted from the HOXC models provided by Schlick's lab at NYU can be found in HOXC_1.tar.gz and HOXC_2.tar.gz.
  There are 6 models: HOXC, Life-like, Life-like-Ac, Life-like-LH, uniformNFR, and uniformNRL. Detailed descriptions of these model can be found in (1).
  The tar files contains 3 representations for each model in xyz file format. Files with only the ".xyz" suffix are representations of the HOXC models. Files with the ".E-R.xyz" suffix were converted from "xyz" to "generalized step parameter" representations and then back to "xyz" using the E-R method. Files with the ".E-A.xyz" suffix converted using the E-A method. The RMSD for each model is shown in Table 1.

**Table 1.** RMSD values associated with converting $C(S)$ representations of HOXC models (1) to "generalized step parameters" and back using Euler-Rodrigues and Euler Angle methods:

| Model | Methods<br>E-R<br>(Å) | <br>E-A<br>(Å) |
|---|---|---|
| HOXC | 0.153 | 0.002 |
| Life-like | 0.154 | 0.002 |
| Life-like-Ac | 0.118 | 0.002 |
| Life-like-LH | 0.050 | 0.002 |
| uniformNFR | 0.059 | 0.003 |
| uniformNRL | 0.028 | 0.003 |

RMSD reported to three significant digits
